# Streptococcal Lancefield polysaccharides are critical cell wall determinants for human group IIA secreted phospholipase A2 to exert its bactericidal effects

**DOI:** 10.1101/269779

**Authors:** Vincent P. van Hensbergen, Elin Movert, Vincent de Maat, Christian Lüchtenborg, Yoann Le Breton, Gérard Lambeau, Christine Payré, Anna Henningham, Victor Nizet, Jos A.G. van Strijp, Britta Brügger, Fredric Carlsson, Kevin S. McIver, Nina M. van Sorge

**Affiliations:** Medical Microbiology, University Medical Center Utrecht, Utrecht, University, Utrecht, The Netherlands; Department of Experimental Medical Science, Section for Immunology and Lund University, Lund, Sweden; Department of Biology, Section for Molecular Cell Biology, Lund University, Lund, Sweden; Heidelberg University Biochemistry Center (BZH), Heidelberg, Germany; Department of Cell Biology & Molecular Genetics and Maryland Pathogen Research Institute, University of Maryland, College Park, Maryland, United States of America; Université Côte d’Azur, CNRS, Institut de Pharmacologie Moléculaire et Cellulaire, Valbonne, France; Department of Pediatrics and University of California, San Diego, La Jolla, CA, United States of America.; Skaggs School of Pharmacy & Pharmaceutical Sciences, University of California, San Diego, La Jolla, CA, United States of America.

## Abstract

Human Group IIA secreted phospholipase A_2_ (hGIIA) is an acute phase protein with bactericidal activity against Gram-positive bacteria. Infection models in hGIIA transgenic mice have suggested the importance of hGIIA as an innate defense mechanism against the human pathogens Group A *Streptococcus* (GAS) and Group B *Streptococcus* (GBS). Compared to other Gram-positive bacteria, GAS is remarkably resistant to hGIIA activity. To identify GAS resistance mechanisms, we exposed a highly saturated GAS M1 transposon library to recombinant human hGIIA and compared relative mutant abundance with library input through transposon-sequencing (Tn-seq). Based on transposon prevalence in the output library, we identified nine genes, including *dltA* and *lytR,* conferring increased hGIIA susceptibility. In addition, seven genes conferred increased hGIIA resistance, which included two genes, *gacH* and *gacI* that are located within the Group A Carbohydrate (GAC) gene cluster. Using GAS 5448 wild-type and the isogenic *gacI* mutant and gacI-complemented strains, we demonstrate that loss of the GAC *N*-acetylglucosamine (GlcNAc) side chain in the *ΔgacI* mutant increases hGIIA resistance approximately 10-fold, a phenotype that is conserved across different GAS serotypes. Increased resistance is associated with delayed penetration of hGIIA through the cell wall. Correspondingly, loss of the Lancefield Group B Carbohydrate (GBC) rendered GBS significantly more resistant to hGIIA-mediated killing. This suggests that the streptococcal Lancefield antigens, which are critical determinants for streptococcal physiology and virulence, are required for the human bactericidal enzyme hGIIA to exert its bactericidal function.

**Author summary:** The human immune system is capable of killing invading bacteria by secreting antimicrobial proteins. Cationic human Group IIA secreted phospholipase A_2_ (hGIIA) is especially effective against Gram-positive bacteria by degrading the bacterial membrane. HGIIA requires binding to negatively charged surface structures before it can penetrate through the thick peptidoglycan layer and reach the target phospholipid membrane. HGIIA is constitutively expressed at high concentrations at sites of possible bacterial entry, e.g. in tears, skin and small intestine. In serum, normal concentrations are low but can increase up to 1,000-fold upon inflammation or infection. *In vitro*, *ex vivo* and *in vivo* experiments suggest an important role for hGIIA in defense against two human pathogens, Group A and Group B *Streptococcus* (GAS, GBS). We demonstrate that the Lancefield cell wall polysaccharides that are expressed by these bacteria, the Group A Carbohydrate (GAC) for GAS and the Group B Carbohydrate (GBC) for GBS, are required for optimal hGIIA bactericidal efficacy by facilitating penetration through the peptidoglycan layer. Given the increased hGIIA resistance of antigen-modified or antigen-deficient streptococci, it will be of interest to determine potential regulatory mechanisms regarding expression of streptococcal Lancefield polysaccharides.

## Introduction

Many important human bacterial pathogens are also common colonizers of mucosal barriers. Occasionally, such pathogens penetrate these physical barriers to invade the underlying tissues and cause infections. Antimicrobial molecules, sometimes also referred to as ‘endogenous antibiotics of the host’, are a critical part of the innate immune response to eradicate these intruders and clear the infection. In humans, one of the most potent bactericidal molecules against Gram-positive bacteria is the secreted enzyme human Group IIA phospholipase A2 (hGIIA) [1,2].

HGIIA belongs to a family of 11-12 secreted phospholipase A_2_ (sPLA_2_) enzymes, which are structurally related and hydrolyze various phospholipids [2–5]. In non-inflamed conditions, hGIIA serum levels are low and not sufficient to kill most Gram-positive bacteria [6]. However, sterile inflammation or infection increases hGIIA expression with concentrations reaching up to 1 µg/ml in serum [7], which is sufficient to kill most Gram-positive pathogens *in vitro*. A unique feature of hGIIA compared to other secreted Phospholipase A_2_ family members is its high cationic charge, which is required for binding to negatively-charged surface structures and for penetration of the thick peptidoglycan layer surrounding Gram-positive bacteria [2,8,9]. The potent bactericidal activity of hGIIA against Gram-positive bacteria has been demonstrated *in vitro*, using recombinant hGIIA, and is suggested by infection experiments that show increased protection from infection using hGIIA transgenic mice [10–16].

To counter the bactericidal effects of hGIIA, pathogens have evolved different resistance mechanisms, for example by suppressing hGIIA expression [17,18] or by increasing the net positive charge of surface structures and membrane. The surface modifications include the addition of positively-charged D-alanine moieties to teichoic acid polymers by the highly conserved *dlt* operon to repulse hGIIA [8] and other cationic antimicrobials [19–22]. In addition, *Staphylococcus aureus* (*S. aureus*) modifies the charge of its bacterial membrane through the molecule MprF [23,24] by adding the cationic amino acid lysine to phosphatidylglycerol (PG), resulting in lysyl-PG [25]. In Group A *Streptococcus* (GAS), the enzyme sortase A (SrtA), a conserved enzyme in Gram-positive bacteria that recognizes proteins with an LPXTG motif and covalently attaches them to peptidoglycan [26,27], was shown to increase hGIIA resistance [12].

Studies with recombinant hGIIA have highlighted differences in intrinsic hGIIA susceptibility between different Gram-positive species, where *Bacillus subtilis* is killed in the low ng/ml concentration range [28,29], and Group A *Streptococcus* (GAS) is one of the most resistant species known to date [12]. Interestingly, this high resistance is not a common trait of streptococcal pathogens since Group B *Streptococcus* (GBS) is killed by concentrations that are approximately 500 times lower compared to those required to kill GAS [11,12]. Streptococci are historically classified by the expression of structurally different Lancefield antigens [30]. Lancefield antigens are cell wall polysaccharides making up approximately 50% of the dry cell wall mass [31]. All GAS serotypes express the Lancefield Group A carbohydrate (GAC), which consists of a polyrhamnose backbone with alternating *N*-acetylglucosamine (GlcNAc) side chains [31], which are important for virulence [32]. In contrast, all GBS serotypes express the Lancefield Group B carbohydrate (GBC), a multi-antennary structure, containing rhamnose, galactose, GlcNAc, glucitol, and significant amounts of phosphate [33]. Both streptococcal species are important human pathogens as they can cause systemic infections associated with high mortality and morbidity [34–36]. Mouse infection models and *ex vivo* studies on human serum from infected patients suggest the importance of hGIIA in defense against lethal infections with GAS and GBS [11,12]. Given the importance of hGIIA in host defense against streptococci, we set out to identify the molecular mechanisms that confer resistance to hGIIA using a comprehensive and unbiased approach.

## Results

### Globally-disseminated M1T1 GAS is highly resistant to hGIIA

A previous study found that GAS strains are among the most resistant Gram-positive bacteria regarding hGIIA-mediated killing [12]. Mutation of *srtA* in the GAS strain JRS4, an *emm6* serotype, increased hGIIA susceptibility by about 50-fold [12]. GAS M1T1 is a globally-disseminated *emm1* clone that is most often responsible for invasive GAS infections in industrialized countries [37,38] and was not included previously in hGIIA studies [12]. GAS strain 5448, a representative M1T1 isolate, showed concentration-dependent killing by recombinant human hGIIA, with an LD_50_ of 0.05 µg/ml (Fig S1). Also, GAS M1T1 resistance mechanisms against hGIIA at least partially overlap with GAS JRS4 *emm6*, since mutation of *srtA* rendered GAS M1T1 approximately 35-fold more susceptible to hGIIA (Fig S1) [12].

### Identification of GAS genes that affect hGIIA susceptibility using Tn-seq

We set out to identify additional genes that affect hGIIA susceptibility of GAS M1T1 using the GAS *Krmit* transposon mutant library [39]. To ensure complete coverage of the library in our experiment, we optimized our hGIIA killing assay to support an inoculum of 10^7^ CFU, using a final concentration of 0.125 µg/ml hGIIA. The Tn-seq experiment with the GAS *Krmit* transposon mutant library consisted of four non-exposed control samples and four hGIIA-treated samples. Each sample contained on average approximately 30 million reads, of which over 90% of the reads aligned once to the GAS M1T1 5448 reference genome (Table S1) [40]. To quantify the number of transposon insertions per gene, we divided the reference genome into 25 nucleotide windows, resulting in 73,182 windows, and mapped each read to a specific window. More than 48% of the windows had at least one read aligned. We identified one gene with an exceptionally high number of transposon insertions at a specific part of the gene (M5005_Spy_1390), suggesting biased insertion of the transposon (Tables S2, S3 and Fig S2). This gene was therefore excluded from further analysis. No other biased transposon insertion sites were observed.

We identified 16 genes that contained a significantly different number of transposon insertions after exposure to hGIIA as indicated by *P*-value of <0.05 (Benjamini-Hochberg (BH) corrected; Fig 1A, Fig S3, and Tables S2-4). Nine of the 16 genes (56%) showed a decrease in transposon insertions compared to untreated controls, indicating that the products of the disrupted genes provide resistance against hGIIA-mediated GAS killing (Fig 1A, Fig S3, Table S3). Three susceptibility genes are located within the *dlt* operon (M5005_Spy_1070, M5005_Spy_1072, M5005_Spy_1073), which is responsible for D-alanylation of teichoic acids [41]. Consistently, this operon was previously linked to GAS resistance against other cationic antimicrobials, such as LL-37 and hGIIA [8,42]. The other six genes with significant fold decrease in transposon insertions are annotated as hypothetical proteins (M5005_Spy_0918 and M5005_Spy_1794), a lactoylglutathione lyase (M5005_Spy_0876), LytR (M5005_Spy_1474) of the LytR/CspA/Psr protein family, the transcriptional regulator FabT (M5005_Spy_1495), and the NAD glycohydrolase inhibitor (M5005_Spy_0140) (Fig S3 and Tables S2, S3).

**Fig 1.**
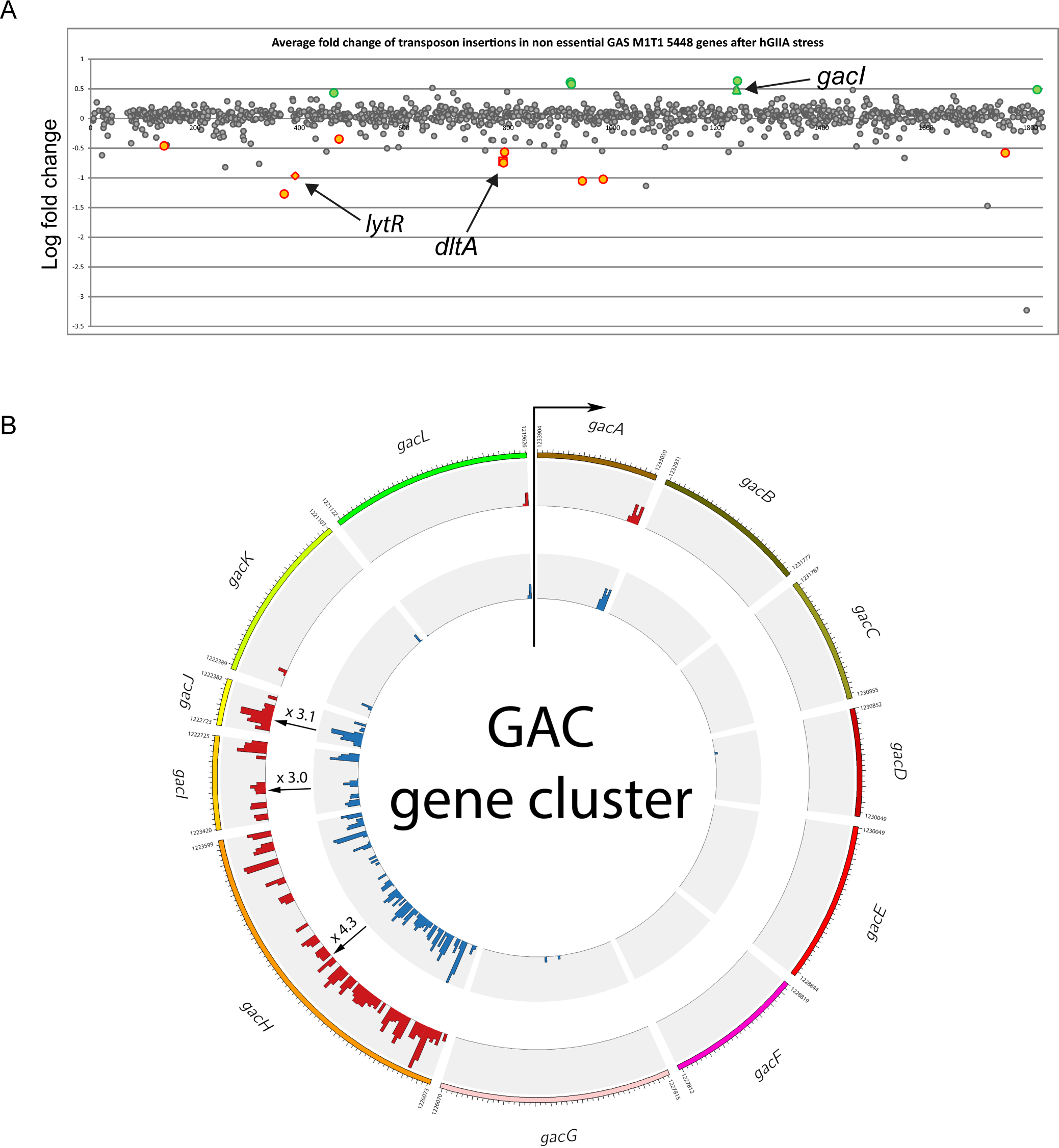
Identifying GAS mutants with different hGIIA susceptibility by Tn-seq analysis. (A) Average log-fold change of transposon insertions in genes of the hGIIA-treated group versus the control group. Grey dots represent genes without significant fold change after hGIIA treatment. Green and orange dots represent genes with significantly increased and reduced transposon insertions after hGIIA exposure, respectively. Significant hits have a calculated BH corrected *p* <0.05. (B) Circos representation of the average transposon insertions of genes within the GAC gene cluster. All genes within this cluster, except for *gacI*, *gacJ*, and *gacH*, were previously identified as essential [39]. For *gacI* and *gacH*, the fold change shown is significant (BH corrected *p* < 0.05), whereas for *gacJ* the fold change is not significant (BH corrected *p* = 0.16).

Seven genes showed a relative increase in the number of transposon insertions after hGIIA exposure, indicating that the products of these genes are important for hGIIA to exert its bactericidal effect (Fig 1A, Fig. S3, Table S4). Five of the six genes (83%) mapped to two gene clusters; one gene cluster is annotated as an ABC transporter (M5005_Spy_0939, M5005_Spy_0940, M5005_Spy_0941) and the other gene cluster is the previously identified 12-gene cluster responsible for biosynthesis of the Group A carbohydrate (GAC) (Fig 1B) [32]. Within the GAC gene cluster, *gacI* and *gacH* (M5005_Spy_0609 and M5005_Spy_0610) showed significantly increased number of transposon insertions. The small downstream gene *gacJ* (M5005_Spy_0611) also demonstrated a 3-fold increase, however, the BH corrected *P*-value is slightly above 0.05. Other genes within the GAC gene cluster are essential or crucial as described previously [39,43]. Finally, *guaB* (M5005_Spy_1857) and the IIC component of a galactose-specific PTS system (M5005_Spy_1399) were identified as their mutation may confer increased resistance to hGIIA (Fig S3 and Tables S2, S4). Overall, the transposon library screen identified genes that confer resistance or are important for the mechanisms of action of hGIIA.

### HGIIA requires the GAC GlcNAc side chain to exert its bactericidal effect against GAS

To validate the Tn-seq findings, we confirmed the involvement of three genes (*dltA*, *lytR*, and *gacI*) by comparing hGIIA-mediated killing of WT GAS with previously generated GAS mutants [32,42,44]. Deletion of *dltA* and *lytR* indeed increased GAS susceptibility to hGIIA-mediated killing by 45-fold and 35-fold, respectively (Fig 2A, B). The *dltA* defect could be restored by re-introduction of the gene on a plasmid (Fig 2A).

**Fig 2.**
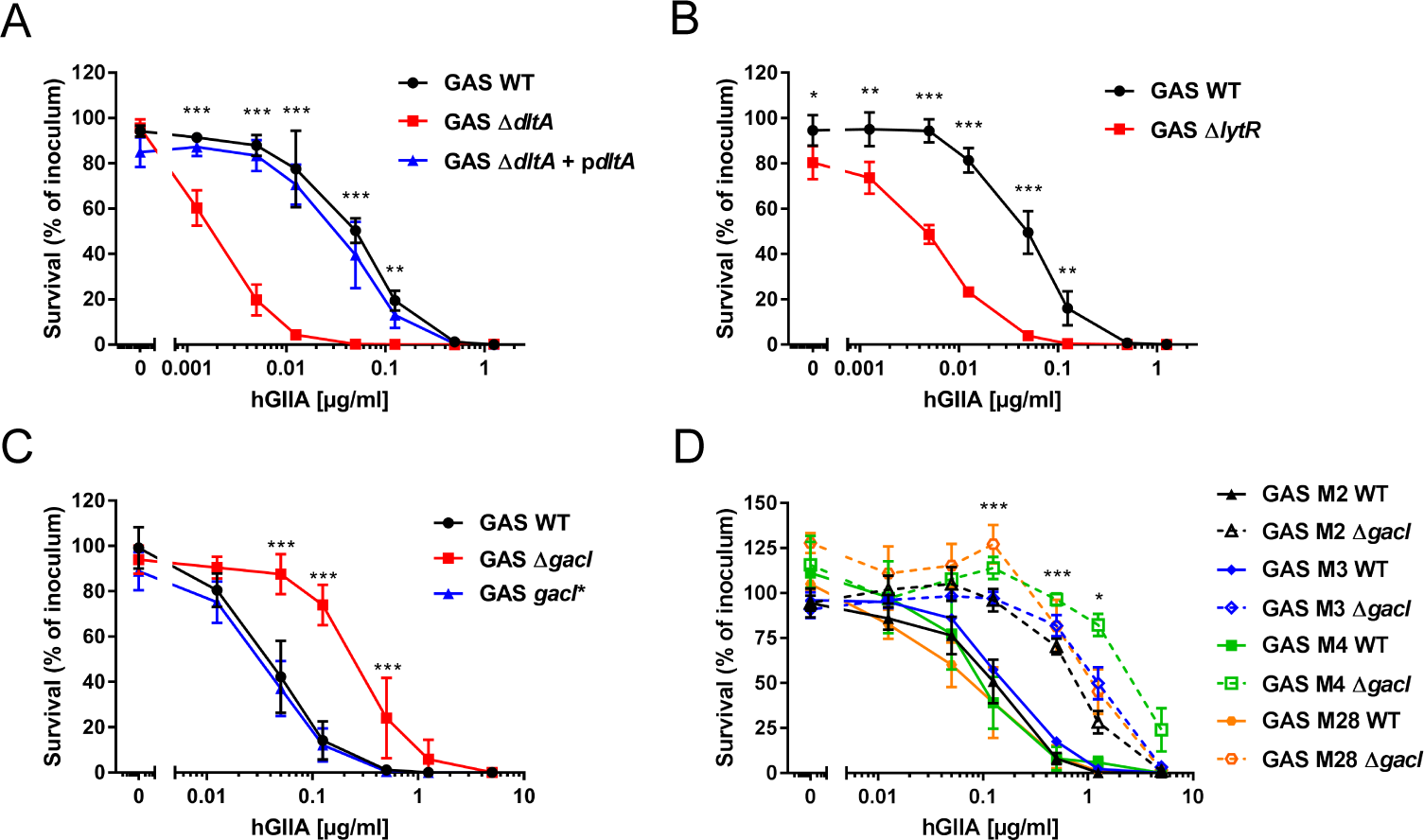
Mutation of *dltA* and *lytR* renders GAS more susceptible to hGIIA, whereas mutation of *gacI* increases hGIIA resistance in multiple GAS serotypes. Deletion of (A) *dltA* or (B) *lytR* increases GAS susceptibility to hGIIA-mediated killing in a concentration-dependent manner. Deletion of *gacI* renders GAS more resistant to hGIIA-mediated killing as shown for (C) 5448 and (D) other tested GAS serotypes. Data represent mean +/- SD of three independent experiments. *, *p* > 0.05; **, *p* > 0.01; ***, *p* > 0.001.

In contrast to *dltA* and *lytR*, mutation of *gacI*, which results in loss of the GAC GlcNAc side chain [45], increased GAS resistance to hGIIA by approximately 10-fold compared to the parental or *gacI*-complemented (*gacI\**) strain (Fig 2C). The GAC is conserved in all GAS serotypes. We therefore questioned whether deletion of *gacI* would have a similar effect on the bactericidal efficacy of hGIIA in four other GAS serotypes (M2, M3, M4, M28). In all serotypes, deletion of *gacI* increased resistance of GAS to hGIIA by 5- to 50-fold (Fig 2D), indicating that hGIIA requires the GAC GlcNAc side chain for optimal bactericidal efficacy in all genetic backgrounds tested.

### Activity and bacterial resistance to hGIIA in human serum

To study the activity of hGIIA in a more physiological setting, we spiked pooled normal human serum with different concentrations of recombinant hGIIA. As described previously [32,46], GAS grows in human serum, a trait that is not influenced by the presence of endogenous hGIIA since addition of the hGIIA-specific inhibitor LY311727 [47] did not affect GAS growth in serum (Fig. S4A). Addition of recombinant hGIIA to human serum potentiated its bactericidal effect compared to the purified assay as reflected by a 5-fold lower LD_50_ (0.01 ug/ml; Fig 3A versus Fig. 2). Interestingly, heat-inactivation of serum reduced the ability of hGIIA to kill GAS by 10-fold compared to active serum, indicating that there are heat-labile factors in serum that potentiate hGIIA efficacy (Fig 3A).

**Fig 3.**
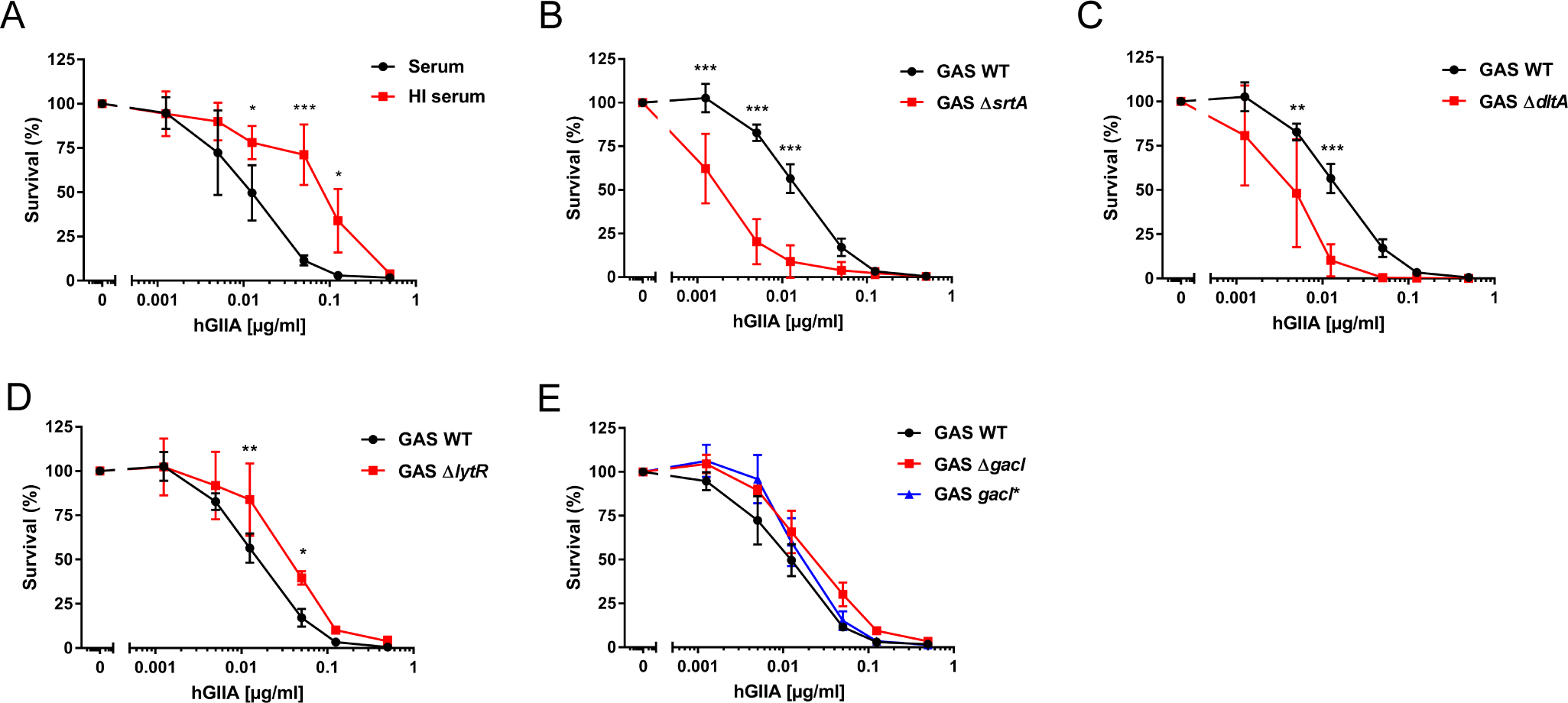
Human serum influences hGIIA efficacy on GAS. (A) A heat-labile factor in serum enhances the ability of hGIIA to kill GAS. The (B) *srtA* and (C) *dltA* GAS mutants retained a susceptible phenotype in hGIIA-spiked serum whereas the (D) *lytR* and (E) *gacI* mutants mutant survive equal to GAS WT under these conditions. Data represent mean +/- SD of three independent experiments. *, *p* > 0.05; **, *p* > 0.01; ***, *p* > 0.001.

We determined how the addition of serum would affect the efficacy of hGIIA to kill the mutants with altered hGIIA susceptibility. We first compared bacterial survival of the WT strain and the individual mutants in normal serum (Fig. S4A). Interestingly, the *lytR* and *srtA* mutant already showed a significant loss of fitness in non-inflamed serum, which is not attributed to the presence of endogenous hGIIA as addition of LY311727 did not impact survival (Fig. S4A). Both *ΔsrtA* and *ΔdltA* bacteria remained more susceptible to hGIIA-mediated killing in serum (Fig. 3B, C), whereas the *ΔlytR* and *ΔgacI* mutants were now equally resistant to WT GAS (Fig. 3D, E). These results reflect the multitude of effects that occur simultaneously in a complex environment such as serum. More specifically, serum likely contains factors that have an opposite effect to hGIIA on *lytR* and *gacI* mutants, such that the net survival of these mutants is equal to WT. Finally, we compared the effect of serum heat-inactivation on hGIIA efficacy in the context of individual mutants (Fig. S4B-D). Similar to WT GAS, heat inactivation of serum reduced the efficacy of hGIIA to kill *ΔsrtA*, *ΔdltA* and *ΔlytR*, suggesting that the hGIIA-potentiating factor(s) is required to kill all mutant in our panel.

### Loss of the GAC GlcNAc side chain delays cell wall translocation of hGIIA

Our observation that GAS *ΔgacI* is more resistant to hGIIA implies that the GAC GlcNAc moiety is important for the function of hGIIA. To assess whether loss of the GAC GlcNAc side chain affected hGIIA binding to bacteria, we first analyzed binding of hGIIA by fluorescence microscopy using a sPLA_2_-specific antibody (Fig. 4A). A visual quantification of hGIIA-stained bacteria indicated reduced binding of hGIIA in the absence of GAC GlcNAc (Fig. 4C). In addition, we observed that the localization of hGIIA on the bacterial surface was affected, where hGIIA predominantly localized to the GAS cell poles in WT bacteria (Fig 4A, B), but distribution became more disperse upon mutation of *gacI* (Fig 4A, B). Since fluorescence microscopy did not allow for more extensive binding assessments, we also quantified binding of recombinant hGIIA to GAS by flow cytometry. At concentrations up to 1 μg/ml, we did not observe differences in hGIIA binding to the three strains (Fig 4D). Only at concentrations of 5 μg/ml, hGIIA showed reduced interaction with the *gacI* mutant compared to GAS WT and *gacI*\*-complemented strains (Fig 4D). The contribution of differential hGIIA binding to GAS is therefore only relevant to specific locations such as in tears which contain up to 30 μg/ml hGIIA [28]. Since hGIIA binding is charge-dependent, we analyzed whether reduced binding at high hGIIA concentrations could be caused by difference in surface charge. Using the highly cationic protein cytochrome C, we indeed observed that the *gacI* mutant has a reduced negative surface charge compared to GAS WT and the *gacI*-* complemented strain (Fig S5), which could likely explain the reduced binding of hGIIA to this mutant.

**Fig 4.**
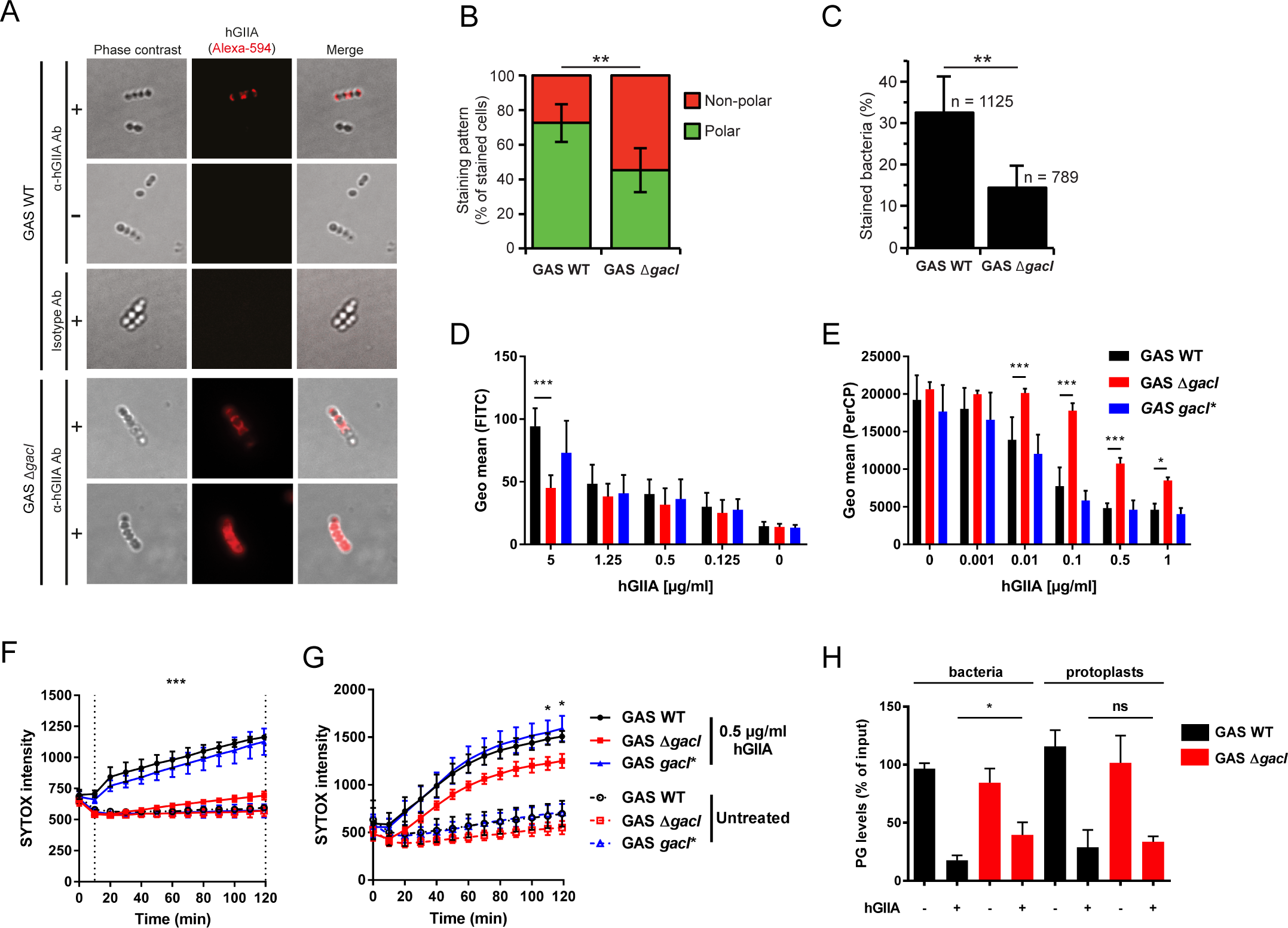
Lack of the GAC GlcNAc side chain delays hGIIA cell wall translocation. (A) Localization of hGIIA (H48Q) on GAS 5448 WT and *ΔgacI* was analyzed by fluorescence microscopy (+) and quantified based on analysis of 10 fields including 307 stained cells from two separate experiments. As control, H48Q hGIIA was omitted (-). HGIIA was detected with a mouse anti-human hGIIA monoclonal antibody or an IgG1 isotype as negative control. Representative bacteria are shown. Quantification of (B) hGIIA localization of hGIIA on GAS WT and GAS *ΔgacI* and (C) the percentage of hGIIA-stained bacteria. (D) Detection of hGIIA binding to GAS 5448 by flow cytometry. (E) Effect of hGIIA on GAS membrane potential after 2 hour incubation. Decreased PerCP signal indicates a disrupted membrane potential. (F) SYTOX green uptake over time by GAS strains or (G) GAS protoplasts after incubation with 0.5 µg/ml recombinant hGIIA. (H) Quantification of PG levels in lipid extracts obtained from WT and *gacI* mutants incubated in the absence or presence of 2 µg/ml hGIIA. Data represent mean +/- SD of at least three independent experiments. ns = not significant, *, *p* > 0.05; **, *p* > 0.01; ***, *p* > 0.001.

Cell wall architecture can significantly affect hGIIA cell wall penetration [2]. To assess how absence of the GAC GlcNAc side chain affected hGIIA cell wall penetration, we measured changes in membrane depolarization over time using the fluorescent voltage-sensitive dye DiOC_2_(3) [48]. In this assay, membrane depolarization results in reduced red fluorescence. HGIIA required at least 5 minutes to penetrate the GAS cell wall since no changes in red fluorescence signal were observed at this time point for any of the strains (Fig S6A). At 30 minutes (Fig S6B), membrane depolarization occurred as visualized by diminished red fluorescence at hGIIA concentrations of 0.1 μg/ml in the GAS WT and the *gacI\**-complemented strain. Compared to these two strains, the *gacI* mutant exhibited limited effects on membrane potential at all time points and all hGIIA measured (Fig 4E and Fig S6). These data suggest that hGIIA reaches the membrane faster in the presence of GAC GlcNAc moieties.

Membrane depolarization likely precedes more pronounced hGIIA-mediated disruption of the membrane that would allow influx of the fluorescent DNA dye SYTOX green, which can only enter damaged membranes [49]. As expected, hGIIA increased the SYTOX signal in GAS WT and GAS *gacI\** in both a time and concentration-dependent manner (Fig 4F and Fig S7A-E). Importantly, addition of LY311727 completely prevented SYTOX influx (Fig. S7F), confirming that our assay indeed reflects hGIIA phospholipase activity on the bacterial membrane. In sharp contrast, SYTOX intensity in GAS *ΔgacI* increased at a much slower rate and never reached the levels of GAS WT and GAS *gacI*\* after two hours. The observed differences in kinetics and severity of hGIIA on membrane depolarization and SYTOX influx in GAS *ΔgacI* compared to GAS WT suggest that the GAC GlcNAc side chain is essential for efficient trafficking of hGIIA through the GAS cell wall.

A recent study demonstrates that GacI is a membrane protein that is required for the intracellular formation of undecaprenyl-P-GlcNAc [45]. Therefore, loss of GacI could alter membrane composition and fluidity to impact the activity of hGIIA on the membrane. To analyze whether phospholipid hydrolysis is affected in GAS *ΔgacI*, we performed the SYTOX influx assay on protoplasts [50]. Unlike the previous SYTOX results with intact bacteria, protoplasts from WT, *ΔgacI* and *gacI*\* strains all became SYTOX positive (Fig 4G and Fig S8), underlining our conclusion that the presence of the cell wall in the *ΔgacI* limits access of hGIIA to the streptococcal membrane. Nonetheless, the significantly lower SYTOX in the *ΔgacI* protoplasts compared to the WT and *gacI*\*-complemented protoplasts (Fig 4G and Fig S8), suggests that the absence of GacI has a minimal impact on hGIIA degradation. To further reinforce this conclusion, we determined the levels of phosphatidylglycerol (PG) in bacteria and protoplasts after treatment with hGIIA (Fig 4H). PG levels were significantly higher in GAS *ΔgacI* after hGIIA treatment compared to WT, whereas equal PG levels were observed in GAS *ΔgacI* and WT after hGIIA treatment (Fig 4H). We therefore conclude that cell wall trafficking and not cell membrane differences are the major determinant of susceptibility differences between GAS WT and *ΔgacI* mutant.

### GBC is important for hGIIA bactericidal activity against GBS

We wondered whether the importance of the GAC for hGIIA activity could be extended to other streptococci such as GBS. As previously described, GBS are generally more sensitive to hGIIA compared to GAS [12]. Indeed, killing of GBS strain NEM316 occurred at substantially lower concentrations of hGIIA compared to GAS M1T1 (compare Fig 5A and 2), also in the presence of serum (Fig. S9). We confirmed that killing depends on the catalytic activity of the enzyme since introduction of an inactivating point mutation (H48Q; Fig 5B) or addition of LY311727 abrogated all killing (Fig 5C). Just as the GAC is the molecular signature for GAS, GBS uniquely express another Lancefield antigen, known as the Group B Carbohydrate (GBC). The GBC is a more complicated structure compared to the GAC and contains significant amounts of phosphate that introduce a negative charge. Unfortunately, there are currently no GBS mutants available with specific structural variations in the GBC. Instead, we assessed the effect of the complete GBC, through deletion of *gbcO* [33], on susceptibility of GBS to hGIIA. Deletion of *gbcO* rendered GBS at least 100-fold more resistant to hGIIA compared to GBS WT (Fig 5A-C), and the phenotype is restored upon complementation with *gbcO* on a plasmid (Fig 5A). We could reproduce the *ΔgbcO* phenotype by treating WT GBS with tunicamycin, an inhibitor of *gbcO*-type transferases (Fig 5D) [33,51]. Finally, as observed in GAS, fluorescence microscopy demonstrated that hGIIA bound to the poles of GBS WT (Fig 5E, F). Unlike to GAS, we did not observe that loss of GBC expression reduced binding of hGIIA at higher concentration of hGIIA as assessed by flow cytometry (Fig S10). In conclusion, these results highlight a key role for streptococcal Lancefield antigens in the bactericidal effect of hGIIA.

**Fig 5.**
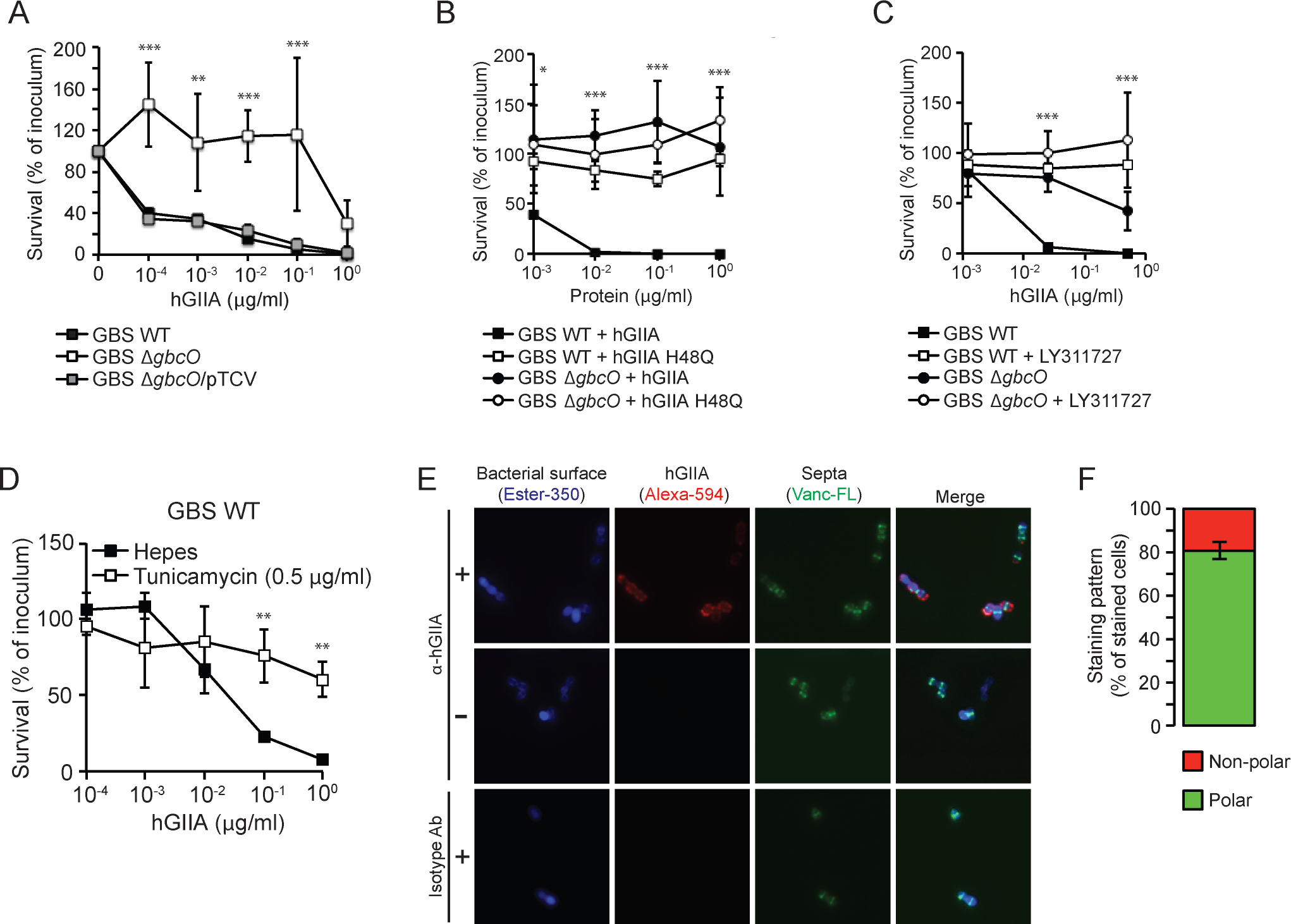
GBS lacking the GBC are resistant to hGIIA. (A) HGIIA kills GBS strain NEM316 WT but not *ΔgbcO* in a concentration-dependent manner and phenotype is restored in complemented strain ΔgbcO/pTCV. The killing is represented as the percentage of CFU surviving after hGIIA exposure compared to the inoculum. GBS killing is prevented when (B) exposed to catalytically inactive hGIIA H48Q and (C) by the hGIIA-specific inhibitor LY311727. (D) Treatment of NEM316 WT with the *gbcO*-type inhibitor tunicamycin reproduces the *ΔgbcO* phenotype with regard to hGIIA-mediated killing GBS more resistant to hGIIA-mediated killing. (E) Visualization of bacteria-bound hGIIA H48Q to GBS NEM316 by fluorescence microscopy (+). As control, H48Q hGIIA mutant protein was omitted (-). hGIIA was detected with a mouse anti-human hGIIA monoclonal antibody. An irrelevant IgG1 isotype antibody served as negative control. The cell wall was labeled with Ester-350, and newly formed septa were visualized with fluorescently labeled vancomycin (Vanc-FL), which stains sites of peptidoglycan insertion. Shown are representative cells. Quantification of hGIIA binding to polar or non-polar regions of GBS are based on analysis of 12 fields including 578 stained cells from two separate experiments. For all other panels, data represent mean +/- SD of three independent experiments. *, *p* > 0.05; **, *p* > 0.01; ***, *p* > 0.001.

## Discussion

Intrinsic resistance to acute phase protein hGIIA varies among Gram-positive bacteria, including among closely-related streptococcal species. GAS, an important cause of lethal infection worldwide, is among the most resistant bacteria, whereas GBS, an important cause of neonatal sepsis and meningitis, is killed by hGIIA at concentrations that are approximately 500-fold lower [12]. For GAS, we confirmed the role of Sortase A and DltA and identified LytR as hGIIA resistance factors. Despite the differences in cell wall composition, i.e. cell wall crosslinking, cell wall associated proteins and membrane physiology, the streptococcal Lancefield antigens are structural requirements for the activity of hGIIA in both GAS and GBS.

HGIIA is approximately 10-fold more effective against GAS when spiked into normal serum compared to heat-inactivated serum, and 5-fold more effective compared to our ‘purified’ system. This corresponds to a previous observation where hGIIA activity was approximately 10-fold greater in serum or plasma than in the protein-depleted serum in studies using *S. aureus* as the target pathogen [52]. This suggests the presence of a heat labile protein in plasma that facilitates hGIIA-mediated killing of Gram-positive bacteria. Heat-inactivation of serum is a well-established method to study the influence of the complement system and also abolishes hGIIA activity in acute phase serum [53]. Since the low basal levels of hGIIA in normal human serum are not sufficient to affect GAS viability, the enhancement could indicate a synergistic effect between hGIIA and the complement system. A recent study shows formation of the Membrane Attack Complex (MAC) on the GAS surface without affecting bacterial viability [54]. It is therefore tempting to speculate that MAC is deposited on Gram-positive bacteria so that bactericidal enzymes, such as hGIIA, can reach the bacterial membrane more easily. Such a cooperative effect between different innate defense mechanisms would not be surprising, since previous studies have already observed that hGIIA synergizes with neutrophil oxygen-dependent mechanisms to kill *Staphylococcus aureus* [55,56]. Finally, the concentrations of hGIIA that are measured in human serum are likely underestimating the true availability of hGIIA since hGIIA attaches to surfaces of blood vessels due to its hydrophobic nature. We speculate that vessel-attached hGIIA may help prevent bacterial dissemination to other tissues, an effect that has not yet been addressed experimentally.

Sortase A, an enzyme that links LPXTG-containing proteins to peptidoglycan, was previously described as a hGIIA resistance factor in GAS serotype M6 [12]. We confirmed that deletion of *srtA* in an GAS M1T1 background similarly sensitizes GAS to hGIIA both in a ‘purified’ as well as a serum environment. Whether a single or multiple LPXTG proteins confer resistance is an unresolved question. Our study suggests that Sortase A-mediated resistance is not caused by a single LPXTG protein since we did not identify a single LPXTG-encoding gene in the Tn-seq screen (Table S5). Possibly, the underlying mechanism is similar to the SrtA-dependent resistance of GAS to the antimicrobial peptide cathelicidin [46], which depends on the accumulation of sorting intermediates at the bacterial membrane. *SrtA* itself was not identified in the transposon library screen since the mutants are lost in the competitive environment likely due to inherent defects in growth [39].

We identified and confirmed a role for the protein LytR in GAS hGIIA resistance. LytR is a member of the LytR-CpsA-Psr (LCP) protein family, a conserved family of cell wall assembly proteins in Gram-positive bacteria [57]. The GAS genome encodes two members of this family, *lytR* (M5005_Spy_1474) and *psr* (M5005_Spy_1099). The fact that we only identified LytR suggests that these proteins have non-redundant, but as yet unidentified, functions. In several Gram-positive pathogens, including *Streptococcus pneumoniae*, *Staphylococcus aureus* and *Bacillus anthracis*, LCP proteins anchor cell wall glycopolymers such as wall teichoic acid (WTA), lipoteichoic acid (LTA) and capsular polysaccharides to the cell envelope and are therefore critical for cell envelope assembly and virulence [57–62]. Additionally, *lytR* homologues in *Bacillus subtilis* and *Streptococcus mutans* contribute to cell wall remodeling by increasing autolysin activity [63,64]. Previously, hGIIA activity has been linked to autolysins; autolysin-deficient mutants are more resistant to hGIIA than their parent strain [65]. A suggested mechanism is that hGIIA displaces positively-charged autolysins from negatively-charged WTA and LTA, resulting in localized peptidoglycan digestion and facilitated movement of hGIIA through the cell wall. Currently, the role of LytR either in GAS cell wall assembly or in the regulation of autolysin activity is not known, but LytR-deficient GAS display altered membrane integrity and potential [66], which could impact hGIIA susceptibility. Moreover, *lytR* has been linked to GAS virulence in two different studies. In the first study, *lytR* mutants in two different GAS M1 backgrounds showed a more virulent phenotype in a subcutaneous murine model of infection, which was suggested to be a result of increased SpeB activity [66]. LytR-mediated regulation of SpeB is unlikely to play a role in hGIIA-mediated resistance in our experiments, since we used washed bacteria. In a more recent study, *lytR* mutants in GAS 5448 M1T1 showed a competitive disadvantage for fitness *in vivo* upon mixed subcutaneous infection [44]. Unfortunately, there is no information regarding pathology or survival of the mice upon infection with the *lytR* mutant added alone [44].

We also identified genes that render GAS more susceptible to hGIIA. *GacH*, *gacI*, and *gacJ* are located in the biosynthesis gene cluster of the GAC, which may suggest that the GAC is a target for hGIIA on the GAS surface. Mutation of *gacI* and *gacJ* results in loss of the GAC GlcNAc side chain [32,45], whereas mutation of *gacH* does not affect side chain formation [32]. We therefore hypothesize that the GAC provides hGIIA resistance through two distinct mechanisms. First, a *gacI*/*J*-dependent mechanism that works through the GAS GlcNAc side chain as important for binding and penetration of hGIIA to the cell membrane. The second mechanism involves GacH but the underlying molecular aspects remain to be determined. The first mechanism seems to conflict with our previous observations that GlcNAc-deficient GAS have decreased virulence capacity due to increased neutrophil killing and increased susceptibility to antimicrobials in serum including LL-37 [32]. However, hGIIA would not have contributed to *in vitro* assays since we used non-inflamed serum or plasma where basal hGIIA concentrations are too low to affect GAS viability [32]. The fact that *gacI* mutants demonstrate reduced survival *in vivo* suggests that the benefits of expressing the GlcNAc side chain outweigh the increased susceptibility to hGIIA. Since GAS already shows high intrinsic resistance towards hGIIA there is no pressure to lose the GlcNAc side chain. It might even be detrimental since it makes GAS more vulnerable to effects of other antimicrobials or yet unidentified host defenses. In contrast to the GAC [31,67], the GBC is a multi-antennary structure and contains anionic charge due to the presence of phosphate [33]. For GBS, the increased hGIIA resistance in GBC-negative *gbcO* mutants is therefore likely explained by the loss of negatively charged groups on the surface. This corresponds to previous observations in *S. aureus*, where loss of the secondary cell wall glycopolymer WTA, increased resistance to several antimicrobial proteins, including hGIIA [10].

Binding of hGIIA to streptococci was reduced when bacteria expressed a modified GAC or lacked complete expression of GBC but these differences were only apparent using high hGIIA concentrations. However, these findings need to be interpreted with caution since possibly only a small portion of the bound hGIIA is required for the bactericidal action of the enzyme. Therefore, even small fluctuations in binding might result in meaningful functional differences. We are currently not able to analyze hGIIA binding at a more sensitive level.

Contrary to our expectations, fluorescence microscopy analysis showed that hGIIA bound to the cell poles of both GAS and GBS. However, the observed binding pattern does not correspond to the reported localization of the GAC and GBC, which are distributed over the entire cell wall as shown by early electron microscopy studies [68,69]. Binding at the septa of dividing bacteria seems a preferred binding site for bactericidal agents due to a high turnover of peptidoglycan which would make penetration easier [70,71]. In addition, the septum is rich in anionic phospholipids [72], a likely target for cationic hGIIA. Finally, the GAS ExPortal, a unique microdomain in the GAS membrane that is enriched in anionic lipids, would be another favored location of binding for the cationic hGIIA [73]. However, the ExPortal is distributed asymmetrically across the GAS surface and not at the cell poles [73]. The fact that we observe a similar binding pattern to GBS and GAS, may indicate that GAS and GBS express a similar protein that localizes at the cell poles and is used by hGIIA as an initial docking site. Importantly, localization became more disperse upon deletion of *gacI* in GAS, possibly suggesting a redistribution of hGIIA-interacting structures. Identification of hGIIA susceptible and resistant GBS mutants using a Tn-seq mutant transposon library may help identify such conserved or homologous hGIIA targets in the GAS and GBS cell wall.

Lack of the GAC GlcNAc side chain most profoundly affected penetration of hGIIA through the cell wall, a mechanism that depends on charge [2,9]. Indeed, membrane depolarization and permeabilization occurs at a much slower rate in the *gacI* mutant compared to WT and complemented strains. This implies that the GAC GlcNAc side chain facilitates penetration of hGIIA through the cell wall in what is referred to as an ‘anionic ladder process’ [2]. Interestingly, the GAC does not contain any charged structures. Therefore, the underlying mechanism may be linked to the previously mentioned autolysin displacement from interaction with the GAC.

In conclusion, we show that the bactericidal agent hGIIA is able to kill GAS in a complex serum environment. However, modification or removal of the Lancefield antigen renders GAS more resistant to the bactericidal activity of hGIIA. Similarly, removing the Lancefield antigen from GBS renders this species also more resistant to the bactericidal activity of hGIIA. The Lancefield antigens, previously thought to be solely involved in physiology, are thus critical cell wall structures for hGIIA to exert its bactericidal effect. The Tn-seq data discussed in this paper provide exciting new insights into the resistance mechanisms of GAS and encourage similar experiments in other streptococci species. Disrupting the resistance mechanisms with therapeutic agents could possibly be sufficient to provide our own immune system the upper hand in clearing invading streptococcal pathogens.

## Materials and Methods

### Bacterial strains and serum

The GAS M1T1 5448 strain was used in this study unless stated otherwise. The 5448*ΔgacI* knockout and *gacI*\* complemented strain [32], the 5448*ΔlytR* [44] and the GAS serotypes M2, M3, M4, and M28 and corresponding *ΔgacI* knockouts [74] were described previously. Preparation and characterization of the GAS M1T1 5448 transposon library was described previously by Le Breton et al., 2015 [39]. All GAS strains were grown in Todd-Hewitt broth (Becton Dickinson) supplemented with 1% yeast extract (Oxoid; THY) as static cultures at 37 °C. Kanamycin (Sigma-Aldrich) was used at a concentration of 300 µg/ml when appropriate. GBS NEM316 WT, *ΔgbcO* and the complemented strains *ΔgbcO*/pTCV were kindly provided by Dr. Mistou [33].

Unless stated otherwise, overnight cultures of GAS were diluted and re-grown to mid-log phase (OD_600nm_ = 0.4), washed and resuspended in HEPES solution (20 mM HEPES, 2 mM Ca^2+^, 1% BSA [pH 7.4]) solution at OD_600nm_ = 0.4 (~1×10^8^ CFU/ml). For GBS strains, overnight cultures of NEM316 WT, *ΔgbcO* and the complemented strains *ΔgbcO*/pTCV were diluted in TH broth and grown to mid-log phase (OD_620nm_ = 0.4 for WT and complemented strains, 0.25 for *ΔgbcO* mutant). Bacteria were then diluted in HEPES solution and pushed rapidly through a 27-gauge needle, a process repeated three times, to disrupt bacterial aggregates. Normal human serum and heat-inactivated serum was obtained from healthy volunteers as described previously [54].

### Identification of GAS resistance determinants against hGIIA

Recombinant hGIIA was produced as described previously [75]. The GAS M1T1 *Krmit* transposon mutant library was grown to mid-log phase in 100 ml THY containing Km and resuspended in HEPES solution to OD_600nm_= 0.4. Four experimental replicates of 100 µl (~ 1×10^7^ CFU) were subsequently incubated in HEPES solution with or without 125 ng/ml hGIIA for 1 hour at 37 °C. After incubation, 3 ml THY was added to all samples and incubated at 37 °C until the mid-log phase was reached (recovery step). Cultures were collected by centrifugation and used for isolation of genomic DNA (gDNA). gDNA was isolated by phenol-chloroform extraction. Samples were barcoded and prepared for Tn-seq sequencing as described previously [76]. Tn-seq sequencing was performed on Illumina NextSeq500 (Sequencing facility University Medical Center, Utrecht, The Netherlands).

Tn-seq data analysis was performed as previously described [76]. In short, barcodes were split using the Galaxy platform [77] and sequences were mapped to the GAS M1T1 5448 genome [40] using Bowtie 2 [78]. The genome was subsequently divided in 25-bp windows and each alignment was sorted and indexed by IGV [79]. Insertions were counted per window and then summed over the genes. Read counts per gene were adjusted to cover only the first 90% of the gene since transposon insertions in the final 10% potentially do not cause a knock-out phenotype. Then, read counts were normalized to the total number of reads that mapped to the genome in each replicate, by calculating the normalized read-count RKPM (Reads Per Kilobase per Million input reads; RKPM = (number of reads mapped to a gene x 10^6^) / (total mapped input reads in the sample x gene length in kbp)). Cyber-T [80] was used to perform statistical analysis on the RKPM values. Genes that contributed to either hGIIA susceptibility or hGIIA resistance were determined when the Benjamini-Hochberg (BH) corrected *p-*value was <0.05. Illumina sequencing reads generated for the Tn-seq analysis were deposited in the European Nucleotide Archive under the accession number PRJEB27626.

### hGIIA susceptibility

Mid-log streptococcal suspensions were diluted 1,000 times in HEPES solution and 10 µl was added to sterile round-bottom 96 well plates (triplicates). Recombinant hGIIA or catalytically-deficient hGIIA mutant enzyme H48Q was serially diluted in HEPES solution or human serum and 10 µl aliquots were added to bacteria-containing wells. For hGIIA inhibition experiments, 50 µM LY311727 was added to the HEPES solution or serum. For GAS, samples were incubated for 2 hours at 37°C, without shaking, PBS was added and samples were 10-fold serially diluted and plated on THY agar plates for quantification. For GBS, bacteria were incubated with hGIIA at 37°C for 30 minutes, the samples were diluted in PBS and plated onto blood agar plates. After overnight incubation 37°C, colony forming units (CFU) were counted to calculate the survival rate (Survival (% of inoculum) = (counted CFU * 100) / CFU count of original inoculum or Survival (%) = (counted CFU * 100) / CFU count at 0 µg/ml hGIIA). For pharmacological inhibition of GBC expression, NEM316 WT bacteria were grown to mid-log phase (OD_620nm_ = 0.4) in the presence of 0.5 mg/ml tunicamycin (Sigma) and used in killing assays as described above.

### Membrane potential and permeability assays

Changes in hGIIA-dependent membrane potential were determined using the membrane potential probe DiOC_2_(3) (PromoKine) [48,81]. Bacterial suspensions (OD_600nm_= 0.4) were diluted 100 times (~1×10^6^ CFU/ml), 100 µl aliquots were divided into eppendorf tubes and incubated with serial dilutions of hGIIA. After incubation at 37°C, 3 mM DiOC_2_(3) was added and incubated at room temperature for 5 minutes in the dark. Changes in green and red fluorescence emissions were analyzed by flow cytometry.

Bacterial staining with the DNA stain SYTOX Green (Invitrogen) is a measurement for membrane permeabilization and an indication of bacterial cell death [49]. Serial dilutions of hGIIA in HEPES solutions were added to wells of a sterile flat-bottom 96 well plate. Bacteria were resuspended in HEPES solution containing 1 µM SYTOX green (OD_600nm_ = 0.4) and added to hGIIA dilutions in a final volume of 100 µl. For hGIIA inhibition experiments, 500 µM LY311727 was added. Fluorescence over time was recorded using FLUOstar OPTIMA (green fluorescence 530 nm emission and excitation 488 nm) at 37°C.

### Surface charge determination

Bacterial surface charge was determined as previously described [81]. Briefly, exponential phase bacteria (OD_600nm_ = 0.4) were washed twice in 20 mM MOPS buffer [pH 7.0] and adjusted to OD_600nm_ = 0.7. After a 10-fold concentration step, 200 µl bacterial aliquots were added to 200 µg cytochrome c (from *Saccharomyces cerevisiae*, Sigma-Aldrich) in a sterile 96-well round-bottom plate. After 10 minutes at room temperature in the dark, the plate was centrifuged, the supernatant was transferred to a sterile 96 well flat-bottom plate and absorbance was recorded at 530 nm. The percentage of bound cytochrome c was calculated using samples containing MOPS buffer only (100% binding) and samples containing MOPS buffer and cytochrome c (0% binding).

### hGIIA surface binding

To determine hGIIA surface binding, 12.5 µl of bacterial cultures in mid-log phase (OD_600nm_ = 0.4 and 0.25 for GBS *ΔgbcO*) were added to wells of a sterile 96-well round-bottom plate (triplicates). hGIIA was serially diluted in HEPES solution without Ca^2+^ and added to the bacteria at indicated concentrations. After 30 minutes incubation at 4°C, bacteria were collected by centrifugation and resuspended in HEPES solution without Ca^2+^ containing 1:300 dilution of anti-sPLA_2_ antibody (Merck Millipore) [28]. After incubation at 4°C for 30 minutes, the samples were washed and incubated with a 1:1,000 dilution of FITC-labeled goat-anti-mouse IgG (SouthernBiotech) or a 1:500 dilution of Alexa Fluor 647 conjugated goat-anti-mouse IgG (Jackson Immuno Research). After washing with HEPES solution without Ca^2+^, samples were fixed with 1% paraformaldehyde and fluorescence was recorded by flow cytometry (FACSVerse, BD Biosciences).

### Fluorescence microscopy

To analyze hGIIA surface localization by microscopy, bacteria were grown in 10 ml broth to mid-log phase and washed with 0.1 M NaHCO_3_ [pH 9]. For GBS, the bacterial septa were stained by addition of a 1:1 mixture of Vancomycin bodipy® FL conjugate (Invitrogen, V34550) and vancomycin (Sigma) at a final concentration of 1.25 µg/ml, during the last generation time of growth. The surface of GBS was stained with Alexa Fluor® 350 Carboxylic acid Succinimidyl ester (Molecular Probes by Life Technologies, A10168) for 1 hour in room temperature. Bacteria were then resuspended in 500 µl HEPES solution and the suspension was divided over two tubes. A final concentration of 10 µg/ml hGIIA H48Q was added to one tube and HEPES solution to the other before a 30 minute incubation at room temperature. The samples were washed and resuspended in 200 µl HEPES solution, then again divided to two tubes. A mouse anti-human hGIIA monoclonal antibody (Clone SCACC353 Cayman Chemical) or an IgG1 isotype control (mouse anti human IgA clone 6E2C1, DAKO) was added to a final concentration of 10 µg/ml to the bacterial suspensions and incubated at RT. After washing, the samples were incubated with 8 µg/ml of Alexa Flour® 594 goat anti-mouse IgG1 (Molecular Probes by Life Technologies, A21125). After 30 min incubation, the samples were washed in HEPES solution and fixed in 4% paraformaldehyde. Ten µl of bacterial suspension were mounted onto microscopic slides (VWR) using MOWIOL (Sigma) mounting medium before viewing the samples using Zeiss Axiovert 200M microscope. Pictures were captured using a 63× objective and the AXIOVISION 4.8 software.

### Hydrolysis of membrane phospholipids

To determine hGIIA efficacy in hydrolyzing membrane phospholipids, the membrane permeabilization assay was modified for protoplasts. Mid-log bacterial suspension were prepared in in protoplast buffer (20% sucrose, 20 mM Tris-HCl, 10 mM MgCl_2_, 2 mM CaCl_2_ [pH 7.4]) containing 1.4 units/µl mutanolysin (Sigma-Aldrich) [50,82,83]. After incubation for 1 hour at 37°C, protoplasts were collected by centrifugation (1,200 rpm 15 minutes) and resuspended in protoplast buffer to an OD_600nm_ = 0.4. Pore formation by hGIIA was monitored using SYTOX Green as described above.

### Quantification of PG levels in lipid extracts

Approximately 3*10^7^ CFU from a mid-log bacterial suspension in HEPES solution, or protoplasts in protoplast buffer, were exposed to 2 µg/ml hGIIA for 30 minutes. Afterwards, bacterial suspensions were centrifuged at 140,000 rpm for 4 minutes and bacterial pellets were resuspended in MeOH. The protoplast suspensions were mixed with MeOH 1:1. Bacterial lipids were extracted under acidic conditions in the presence of 10 pmol PG standards (PG 14:1/14:1, PG 20:1/20:1 and PG 22:1/22:1) as described [84]. Lipid extracts were resuspended in 60 µl methanol and diluted 1:10 in 96 wells plates (Eppendorf twintec 96, colorless, Sigma, Z651400-25A) prior to measurement. Measurements were performed in 10 mM ammonium acetate in methanol. Samples were analyzed on an AB SCIEX QTRAP 6500+ mass spectrometer (Sciex, Canada) with chip-based (HD-D ESI Chip, Advion Biosciences, USA) electrospray infusion and ionization via a Triversa Nanomate (Advion Biosciences, Ithaca, USA) as described [84]. PG species were measured by neutral loss scanning selecting for neutral loss of m/z 189. Data evaluation was done using LipidView (ABSciex).

### Statistical analysis

GraphPad Prism 6 was used to perform statistical analysis. An unpaired two-tailed Student’s *t*-test was used to compare the means of two groups. A 2-way ANOVA with Bonferroni multiple comparison test was used to compare multiple groups. Data shown are mean ± SD.

## Acknowledgements

The authors would like to thank Dr. Michel-Yves Mistou (INRA, Jouy-en-Josas, France) for providing his *gbcO* mutant and complemented GBS strains.

